# Optimization of long-term human iPSC-derived spinal motor neuron culture using a dendritic polyglycerol amine-based substrate

**DOI:** 10.1101/2021.09.14.460098

**Authors:** Louise Thiry, Jean-Pierre Clément, Rainer Haag, Timothy E Kennedy, Stefano Stifani

**Affiliations:** Department of Neurology and Neurosurgery, Montreal Neurological Institute-Hospital, McGill University, 3801, rue University, Montreal (Quebec) H3A 2B4, Canada; Institute of Chemistry and Biochemistry, Freie Universität Berlin, Takustrasse 3, Berlin, 14195, Germany

**Keywords:** Human iPSCs, spinal motor neurons, dendritic polyglycerol amine, multi-electrode array, single cell RNA sequencing

## Abstract

Human induced pluripotent stem cells (h-iPSCs) derived from healthy and diseased individuals can give rise to many cell types, facilitating the study of mechanisms of development, human disease modeling, and early drug target validation. In this context, experimental model systems based on h-iPSC-derived motor neurons (MNs) have been used to study MN diseases such as spinal muscular atrophy and amyotrophic lateral sclerosis. Modeling MN disease using h-iPSC-based approaches requires culture conditions that can recapitulate in a dish the events underlying differentiation, maturation, aging, and death of MNs. Current h-iPSC-derived MN-based applications are often hampered by limitations in our ability to monitor MN morphology, survival, and other functional properties over a prolonged timeframe, underscoring the need for improved long-term culture conditions. Here we describe a cytocompatible dendritic polyglycerol amine (dPGA) substrate-based method for prolonged culture of h-iPSC-derived MNs. We provide evidence that MNs cultured on dPGA-coated dishes are more amenable to long-term study of cell viability, molecular identity, and spontaneous network electrophysiological activity. The present study has the potential to improve iPSC-based studies of human MN biology and disease.

## Introduction

A major goal when studying human neurodegenerative diseases is to generate, culture, and study human neuronal cells displaying disease-relevant phenotypes. The use of human induced pluripotent stem cell (h-iPSC)-derived neurons has provided the opportunity to model a variety of neurodegenerative diseases, thus facilitating the development of new therapeutics. For instance, loss of motor neuron (MN) functions is associated with several devastating neurodegenerative diseases including spinal muscular atrophy and amyotrophic lateral sclerosis (ALS) (Kanning *et al.*, 2010; Nijssen *et al.*, 2017). Several protocols exist to obtain MNs from h-iPSCs (*e.g.*, Li *et al.*, 2005; Qu *et al.*, 2014; Amoroso *et al.*, 2013; Du *et al.*, 2015), and these cells have been used to study the pathophysiology of MNs from ALS and spinal muscular atrophy patients (*e.g.*, Ebert *et al.*, 2009; Egawa *et al.*, 2012; Chen *et al.*, 2014; Kiskinis *et al.*, 2014).

An important goal of MN disease modeling using h-iPSCs is to implement culture conditions that can faithfully recapitulate in a dish the processes underlying MN degeneration. This objective requires the ability to monitor molecular and functional properties of MNs, such as gene/protein expression, morphology, and electrical activity, in long-term cultures of MNs “aged” *in vitro*. However, prolonged culture of h-iPSC-derived MNs is challenging with most commonly used MN derivation protocols. Most notably, h-iPSC-derived MNs typically coalesce into large cell aggregates, and eventually detach from their substrate after continued culture, thus making long-term studies technically challenging (Kuijlaars et al. 2016; Taga et al. 2019; Thiry et al. 2020). Studies relying on endpoint assays requiring immunocytochemical staining, electrophysiological recordings of networks with multi-electrode array (MEA) devices, or single cell RNA sequencing (scRNAseq) applications are hindered by the presence of these large MN clusters, highlighting a critical need to develop improved h-iPSC-derived MN culture conditions that would address these limitations and facilitate long-term studies. Here we describe the use of a dendritic polyglycerol amine (dPGA) (Hellmund *et al.*, 2014) as a new coating substrate providing improved cytocompatible conditions for long-term cultures of h-iPSC-derived MNs, thus allowing continued evaluation of cell viability, molecular identity, and spontaneous network electrophysiological activity. These culture conditions also facilitate scRNAseq studies of mature MNs due to decreased proportions of damaged cells when compared to more conventional coating protocols. These results demonstrate the value of using a dendritic polyglycerol amine substrate for real-time, long-term qualitative and quantitative analysis of h-iPSC-derived MN cultures.

## Materials and methods

### Generation of motor neurons from human iPSCs

Human iPSC line NCRM-1 (male) was obtained from the National Institutes of Health Stem Cell Resource (Bethesda, MD, USA). Cells were differentiated into neural progenitor cells (NPCs), and subsequently MN progenitor cells (MNPCs) and MNs, as described previously (Thiry et al., 2020). After 24 days of differentiation, MNPCs were dissociated and plated at 50,000 cells/well on coverslips coated only with Matrigel (Thermo-Fisher Scientific; Cat. No. 08-774-552) or with dendritic polyglycerol amine (dPGA: ~95 kDa, 50% amine content) (Hellmund et al 2014) plus Matrigel. For the latter condition, coverslips were coated with 300 μL/coverslip of a solution of PBS with dPGA at varying concentrations (10, 25 or 50 μg/mL), and returned to the incubator for at least one hour before rinsing with PBS and coating with Matrigel on top of dPGA. MNs were differentiated in a chemically defined neural medium including DMEM/F12 supplemented with GlutaMAX (1/1; Thermo-Fisher Scientific; Cat. No. 35050-061), Neurobasal medium (1/1; Thermo-Fisher Scientific; Cat. No. 21103-049), N2 (0.5x; Thermo-Fisher Scientific; Cat. No. 17504-044), B27 (0.5x; Thermo-Fisher Scientific; Cat. No. 17502-048), ascorbic acid (100 μM; Sigma-Aldrich; Cat. No. A5960), antibiotic–antimycotic (1x; Thermo-Fisher Scientific; Cat. No. 15240-062), 0.5 μM retinoic acid, 0.1 μM purmorphamine, 0.1 μM Compound E (Calbiochem; Cat. No. 565790), insulin-like growth factor 1 (10 ng/mL; R&D Systems; Minneapolis, MN; Cat. No. 291-G1-200), brain-derived neurotrophic factor (10 ng/mL; Thermo-Fisher Scientific; Cat. No. PHC7074) and ciliary neurotrophic factor (10 ng/mL; R&D Systems; Cat. No. 257-NT-050). The MN culture medium was replaced every other day for 6 days and the MN cultures were characterized by immunocytochemistry as described below. For MEA recording, MNs were dissociated, plated at ~50,000 cells/well on Matrigel-coated or dPGA/Matrigel-coated wells of a CytoView MEA 24-well plate (Axion Biosystems; Atlanta, GA, USA; Cat. No. M384-tMEA-24W), and cultured with the same MN medium described above. Culture medium was replaced every three days for one month, during which cells were recorded every week as described below.

### Characterization of human iPSC-derived cells by immunocytochemistry

Immunocytochemical characterization of induced MNs was performed as described previously (Methot *et al.*, 2018). The following primary antibodies were used: mouse anti-HOMEOBOX PROTEIN HB9 (HB9) (1/30; DSHB; Iowa City, IA, USA; Cat. No. 81.5C10-c); mouse anti-ISLET1 (ISL1) (1/30; DSHB; Cat. No. 39.4D5-c); rabbit anti-LIMB HOMEOBOX CONTAINING 3 (LHX3) (1/100; Abcam; Toronto, ON, Canada; Cat. No. ab14555); goat anti-FORKHEAD BOX PROTEIN 1 (FOXP1) (1/100; R&D Systems; Cat. No. AF4534); and mouse anti-neurofilament protein (2H3) (1/35; DSHB; Cat. No. 2H3-c). The indicated DSHB monoclonal antibodies were obtained from the Developmental Studies Hybridoma Bank, created by the NICHD of the NIH and maintained at The University of Iowa, Department of Biology, Iowa City, IA 52242. Secondary antibodies against primary reagents raised in various species were conjugated to Alexa Fluor 488, Alexa Fluor 555, or Alexa Fluor 647 (1/1,000, Invitrogen; Burlington, ON, Canada). For quantification, images were acquired using an Axio Observer Z1 microscope connected to an AxioCam camera and using ZEN software (Zeiss). For each culture and each condition, images of >500 cells in 3 random fields were taken with a 20X objective and analyzed with Image J.

### Multi-electrode array recording of human iPSC-derived motor neuron cultures

After 24 days of differentiation, h-iPSC-derived MNs were plated at ~50,000 cells/well on Matrigel-coated or dPGA/Matrigel-coated wells of CytoView MEA 24-well plates (Axion Biosystems). Recordings from 16 electrodes per well were conducted using a Maestro (Axion BioSystems) MEA recording amplifier with a head stage maintaining a temperature of 32°C at 5% CO2. MEA plates were allowed to equilibrate for ∼3 min prior to the 5 min recording of spontaneous activity. Data were sampled at 12.5 kHz, digitized, and analyzed using Axion Integrated Studio software (Axion BioSystems) with a band-pass filter (200–5000 Hz). The spontaneous electrophysiological activity of the h-iPSC-derived cells was recorded every 7 days post-plating for 3 weeks: at day-32 (one-week post-plating), day-39 (two weeks post-plating), and day-46 (three weeks post-plating). The following electrophysiological parameters were analyzed: percentage of active electrodes, spike frequency, and firing rate.

### Preparation of single-cells suspensions

Solutions of single-cells in suspension were prepared from MNs cultured for 39 days (day 0 defined as start of NPC induction), approximately 2 weeks post-plating on either Matrigel only or dPGA (50 μg/mL)/Matrigel, as previously described (Thiry et al., 2020). Cells were counted to evaluate cell concentration and viability before scRNAseq.

### Single cell RNA sequencing

Cells were sequenced on a single-cell level using the microdroplet-based protocol 10X Genomics Chromium Single Cell 3’ Solution, followed by Illumina sequencing at the McGill and Genome Quebec Innovation Center (https://cesgq.com/en-services). The 10X cDNA libraries were sequenced at a depth of 50,000 reads per cell on a HiSeq4000 system (Illumina).

### In silico analysis

The raw scRNAseq data were first processed using the 10X pipeline (Cell Ranger) to demultiplex and align the sequences to the version GRCh38 of the human genome. The gene/cell matrices produced by Cell Ranger were used in the quality control processing and analysis. All the subsequent data analysis was conducted in R version 3.5.2 using Bioconductor Software Version 3.8. All packages are publicly available at the Bioconductor project (http://bioconductor.org). The workflow used for the analysis was adapted from a workflow previously developed to characterize h-iPSC-derived MNs (Thiry et al., 2020) adapted from a published workflow designed to accommodate droplet-based systems such as 10X Chromium (Lun et al. 2016).

### Statistics

To detect differences in the proportion of total MNs, lateral motor column (LMC) MNs, or median motor column (MMC) MNs according to the substrate used for coating (i.e.; Matrigel alone or Matrigel in combination with either 10 μg/mL dPGA, 25 g/mL dPGA, or 50 μg/mL dPGA), nonparametric Friedman test and Dunn’s multiple comparisons post hoc test were used, because the variables did not fit a normal distribution (assessed by Kolmogorov-Smirnov test). Kruskal-Wallis and Dunn’s multiple comparisons post hoc test was used to detect differences in the number of single-cells remaining attached to the coverslips after one-week post-plating, according to the substrate used for coating (same as above). Two-way ANOVA and Tukey’s multiple comparison post hoc test were used to detect differences in the percentages of single-cells remaining attached to the coverslips at different time-points (one-, two- and three-weeks post-plating) according to the substrate used for the coating (i.e.; Matrigel alone or Matrigel and either 10 μg/mL dPGA, 25 g/mL dPGA, or 50 μg/mL dPGA). The statistical significance was set at p <0.05.

## Results

### Differentiation of human iPSC-derived motor neurons cultured on dPGA/Matrigel

Spinal MNPCs and MNs were generated from the h-iPSC NCRM-1 line as described previously (Thiry et al., 2020). After 25 days of differentiation using standard culture conditions, 64±6% of induced cells expressed the typical pan-MN markers ISL1 and HB9 (Amoroso *et al.*, 2013) (Figure 1A and 1C; “Matrigel”). A large proportion of the h-iPSC-derived ISL1^+^/HB9^+^ MNs (48±9%) expressed FOXP1, a known marker of limb innervating MNs of the LMC (Dasen et al. 2003 and 2008; Amoroso et al. 2013) and did not express LHX3 (Thaler et al. 2002; Agalliu et al. 2009), thus exhibiting the typical LMC MN molecular profile (FOXP1^+^/LHX3^−^) (Figure 1B; top row; “Matrigel”; and Figure 1C; “LMC”). In contrast, only 14±6% of the h-iPSC-derived MNs were FOXP1^−^ and expressed LHX3, corresponding to the molecular profile (LHX3^+^/FOXP1^−^) of MNs of the MMC (Figure 1B; top row; “Matrigel”; and Figure 1C; “MMC”).

**Figure 1:**
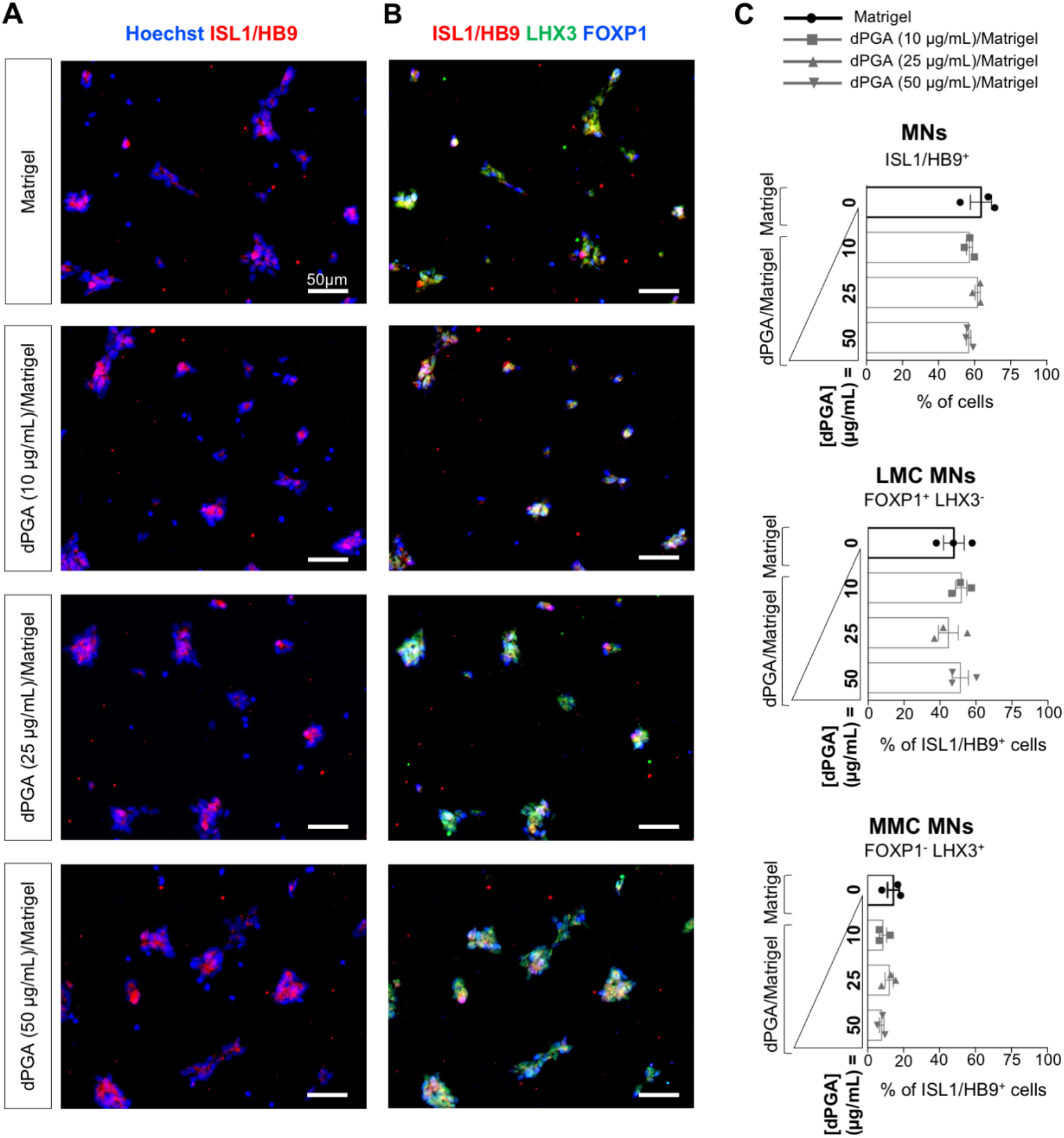
The dPGA/Matrigel substrate does not affect the differentiation of h-iPSCs into spinal motor neurons. **A-B)** Representative images of MNs plated on Matrigel only vs dPGA (10, 25 or 50 μg/mL) and Matrigel combined, and co-stained with the known pan-MN marker HB9/ISL1 **(A)**, as well as anti-LHX3 and anti-FOXP1 antibodies **(B)** after 25 days of culture. **C)** Quantification of all MNs identified as ISL1^+^/HB9^+^, LMC MNs identified as ISL1^+^/HB9^+^/FOXP1^+^/LHX3^−^ and MMC MNs identified as ISL1^+^/HB9^+^/LHX3^+^/FOXP1^−^. Scale bars, 50 μm. n = 3 cultures/condition (with >500 cells in 3 random fields for each culture). Friedman test and Dunn’s multiple comparisons post hoc test. All p values are > 0.05, i.e., not statistically significant. Data are shown as mean ± SEM. MMC, medial motor column; LMC, lateral motor column.

Our work and that of others has shown that h-iPSC derived MNs usually coalesce into large cell clusters which gradually detach from the culture device (Kuijlaars et al. 2016; Taga et al. 2019; Thiry et al. 2020). This situation is most likely due to the degradation of the protein-based substrate (Matrigel or laminin) commonly used for coating, a process that can mediated by enzymes secreted by the cells (reviewed in Schmidt et al 2018; Clement et al. submitted). To address this shortcoming, we sought to test the effect of a non-peptide polymer substrate, dPGA, that was shown to improve primary neuron culture conditions, possibly due to its resistance to degradation by cell-secreted proteases (Clement et al. submitted).

We first asked whether the use of the dPGA compound as a coating substrate in combination with Matrigel would affect the differentiation of h-iPSCs into MNs. MNs were plated on either Matrigel-coated or dPGA/Matrigel-coated coverslips, with varying concentrations of dPGA (10, 25 or 50 μg/mL), followed by analysis by immunocytochemistry after 25 days of culture (Figure 1). A similar proportion (~60%) of the cells plated on Matrigel only or dPGA/Matrigel expressed the pan-MN markers ISL1 and HB9 (Figure 1C; MNs: 64±6% for Matrigel, 57±3% for dPGA10, 62±3% for dPGA25 and 57±2% for dPGA50; not statistically significant, Friedman test and Dunn’s multiple comparisons *post hoc* test). Furthermore, no statistically significant difference was found in the proportion of FOXP1^+^/LHX3^−^ LMC MNs and FOXP1^−^/LHX3^+^ MMC MNs present in the cultures grown on either Matrigel only or dPGA/Matrigel, regardless of the concentration of dPGA used (Figure 1C; LMC MNs: 48±9% for Matrigel, 52±5% for dPGA10, 45±9% for dPGA25 and 51±8% for dPGA50; MMC MNs: 14±6% for Matrigel, 9±4% for dPGA10, 12±4% for dPGA25 and 8±3% for dPGA50; p>0.05, Friedman test and Dunn’s multiple comparisons post hoc test). Together, these observations show that the dendritic polyglycerol amine coating does not detectably interfere with the differentiation of h-iPSCs into MNs, suggesting that it can be used for the culture of h-iPSC derived MNs.

### Improved long-term culture of human iPSC-derived motor neurons on dPGA/Matrigel

To test whether a dendritic polyglycerol amine coating would enhance long-term culture of h-iPSC-derived MNs, we analyzed induced cultures at later time points. As early as one-week post-plating on Matrigel only (i.e.; 32 days (D32) after the start of the differentiation protocol), h-iPSC-derived MNs coalesced into large cell clusters, with their cell bodies crowded into clumps and their processes hyper-fasciculated into well-defined bundles (Figure 2A and 2E). In contrast, at the same time-point, h-iPSC-derived MNs plated on dPGA/Matrigel remained more homogeneously distributed over the coverslip, with their processes creating widespread arborizations, regardless of the three tested dPGA concentrations (Figure 2B-D and 2F). The number of cells identified as single-cells remaining attached to the coverslip was systematically increased by the use of the dPGA substrate (Figure 2G). Interestingly, this effect appeared to be proportional to the concentration of dPGA used, with 10 μg/mL leading to a 4.7±2.7-fold increase, 25 μg/mL leading to a 5.3±1.8-fold increase and 50 μg/mL leading to a 7±2-fold increase in the number of single-cells. Only the use of dPGA at 50 μg/mL led to a statistically significant effect, suggesting that this concentration is the most effective to improve h-iPSC-derived MN culture conditions.

**Figure 2:**
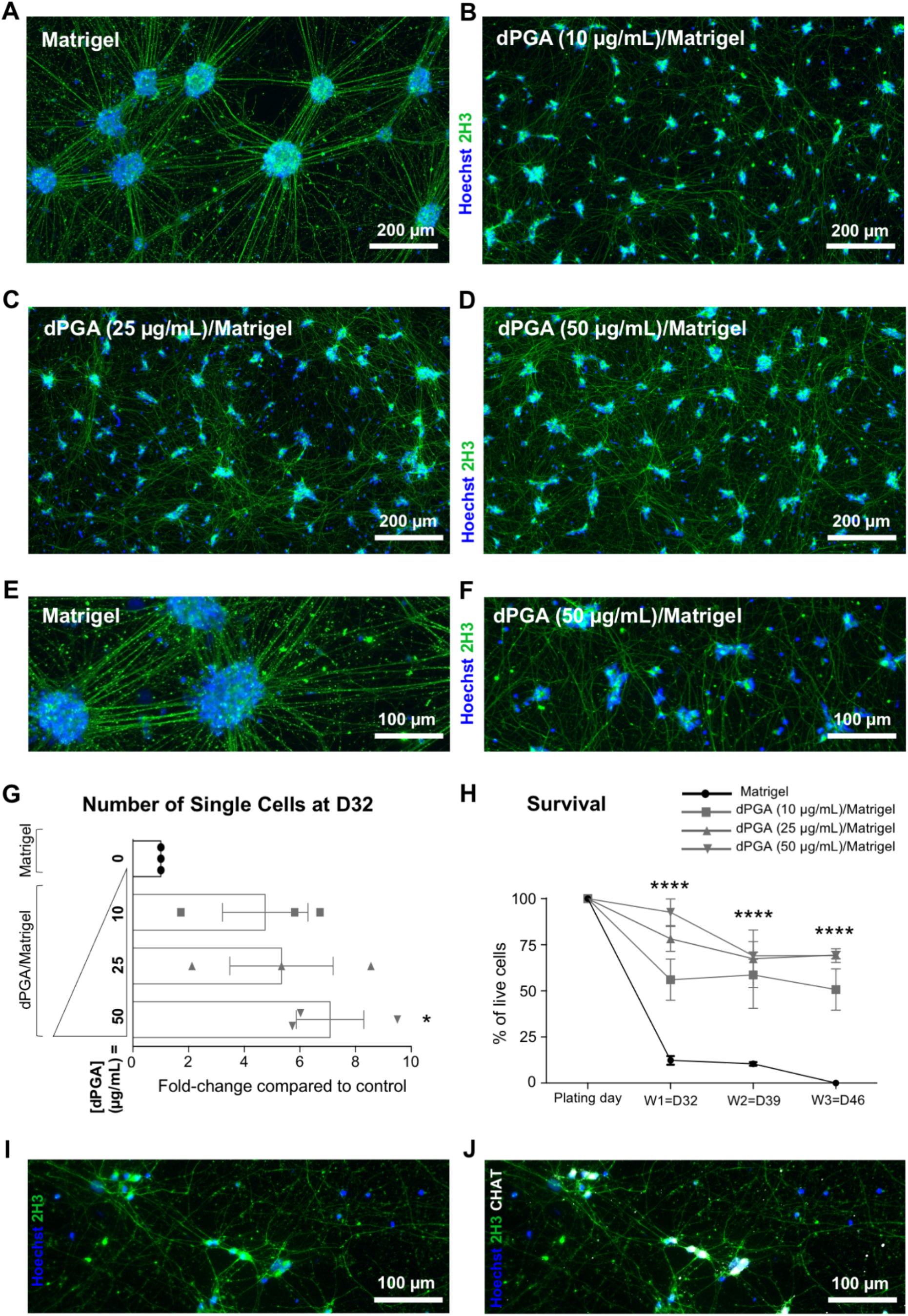
The dPGA/Matrigel substrate enhances long-term culture of human iPSC-derived motor neurons. **A-D)** Representative images of MNs one-week post-plating (i.e., 32 days of differentiation) on either Matrigel only **(A)**, dPGA (10 μg/mL)/Matrigel **(B)**, dPGA (25 μg/mL)/Matrigel **(C)**, or dPGA (50 μg/mL)/Matrigel **(D)**, and stained with Hoechst (blue) and the anti-neurofilament 2H3 antibody (green). Scale bars, 200 μm. **E-F)** Close-up views of MNs plated on Matrigel only **(E)** vs dPGA (50 μg/mL)/Matrigel **(F)**. Scale bars, 100 μm. **G)** Fold change of the number of single cells counted one-week post-plating (i.e., 32 days of differentiation) on either Matrigel only (black) or dPGA/Matrigel (grey). n=3 cultures/condition (with 3 random fields for each culture); *, p<0.05; Kruskal-Wallis and Dunn’s multiple comparisons post hoc test. **H)** Cell number curves from cultures plated on Matrigel only vs dPGA/Matrigel showing the percentages of cells remaining attached to the coverslip after one (i.e., 32 days of differentiation), two (i.e., 39 days) and three (i.e., 46 days) weeks post-plating, as compared to the number of cells plated (100%) (n=5 cultures/condition). Both factors (substrate and time) significantly affected cell adherence to the coverslips (p<0.0001; Two-way ANOVA), and Tukey’s multiple comparisons post hoc tests revealed a statistically significant improvement with dPGA (10, 25, or 50 g/mL)/Matrigel compared to Matrigel only (****, p<0.0001). Furthermore, dPGA (25 and 50 μg/mL) were statistically significantly more efficient then dPGA (10 μg/mL) (p<0.05), while dPGA (25 μg/mL) vs dPGA (50 μg/mL) was not significantly different (Tukey’s multiple comparisons post hoc tests). Data are shown as mean ± SEM. **I-J)** Representative images of MNs two-month post-plating (i.e., 81 days of differentiation) on dPGA (50 μg/mL)/Matrigel and stained with Hoechst (blue) and anti-neurofilament antibody 2H3 (green) **(I)** or Hoechst (blue), anti-neurofilament antibody 2H3 (green) and MN marker anti-CHAT antibody (white) **(J)**. Scale bars, 100 μm.

To evaluate whether this effect of the dendritic polyglycerol amine coating was stable over time, we next compared the percentages of cells remaining attached to Matrigel- or dPGA/Matrigel-coated coverslips after one week, two weeks and three weeks post-plating (Figure 2H). Both the type of coating substrate (e.g., Matrigel only vs dPGA/Matrigel) and the culture time significantly affected cell adherence to the coverslips. As early as one-week post-plating, less than 10% of the cells plated on Matrigel-coated coverslips remained attached and this number decreased over time with all cells becoming detached at three weeks post-plating (Figure 2H, black line). In contrast, a minimum of 50% of the cells plated on dPGA/Matrigel-coated coverslips remained attached over-time, even three weeks after plating (Figure 2H, grey lines) (p<0.0001; Two-way ANOVA). This effect was proportional to the concentration of dPGA used, although each of the three tested concentrations led to a statistically significant increased adherence when compared to Matrigel only (****, p<0.0001 for dPGA (10, 25, or 50 g/mL)/Matrigel compared to Matrigel only, Tukey’s multiple comparisons post hoc tests). A dPGA concentration of 10 μg/mL was significantly less efficient than 25 or 50 μg /mL (p<0.05 for dPGA (10 μg/mL) vs dPGA (25 or 50 μg/mL; Tukey’s multiple comparisons tests), and there was no statistical difference between the effect obtained with a dPGA concentration of 25 or 50 μg/mL. As shown in Figure 2I-J, MNs grown on dPGA (50 μg/mL)/Matrigel could be maintained in culture for at least 2-month post-plating without clumping, and they remained scattered over the coverslip and created widespread arborizations with their processes. Together, these results provide evidence that using the dPGA substrate in conjunction with Matrigel improves the long-term culture conditions of h-iPSC-derived MNs, allowing real-time, long-term qualitative and quantitative analysis of these cells.

### Long-term multielectrode array recordings of human iPSC-derived motor neurons cultured on dPGA/Matrigel

Electrophysiological characterization of h-iPSC-derived MNs is crucial to assess their function. Multielectrode arrays are particularly suited for this purpose as they enable the recording of large neuronal populations through the detection of extracellular voltages. As reported by others (Kuijlaars et al. 2016; Taga et al. 2019), we observed that h-iPSC-derived MNs plated on Matrigel-coated MEA wells formed large cell aggregates (Figure 3A). This made it difficult to record the electrophysiological activity of these cells, as reflected by a low number of electrodes with persistent activity during the three-week recording period (Figure 3C, black line). To test whether the use of dPGA/Matrigel coating could improve MEA analysis of h-iPSC-derived MNs, we recorded h-iPSC-derived MNs plated on either Matrigel- or dPGA/Matrigel-coated MEA plates, with varying concentrations of dPGA (10, 25 or 50 μg/mL), over a three-week period after plating (Figure 3). In contrast to the h-iPSC-derived MNs plated on Matrigel alone, which coalesced into large cell clusters as early as one-week post-plating (Figure 3A), h-iPSC-derived MNs plated on dPGA/Matrigel remained evenly distributed on the MEA wells (Figure 3B). As a result, each of the 16 electrodes in each of the dPGA/Matrigel-coated wells remained in the immediate vicinity of a number of cells for the three-week recording periods. This observation was reflected by a statistically significant increase in the number of electrodes with persistent activity over these recording periods in dPGA/Matrigel-coated wells (Figure 3C, grey lines) as compared to Matrigel-coated wells (Figure 3C, black line). This effect was consistent across the three tested concentrations of the dPGA substrate, without statistical difference between each concentration. Consequently, over the three-week recording periods, both the number of spikes (Figure 3D) and the mean firing rate (Figure 3E) were statistically significantly increased in dPGA/Matrigel-coated wells (grey lines) as compared to Matrigel-coated ones (black line). Together, these results suggest that using the dPGA substrate in combination with Matrigel enables stable long-term MEA recordings of h-iPSC-derived MNs.

**Figure 3:**
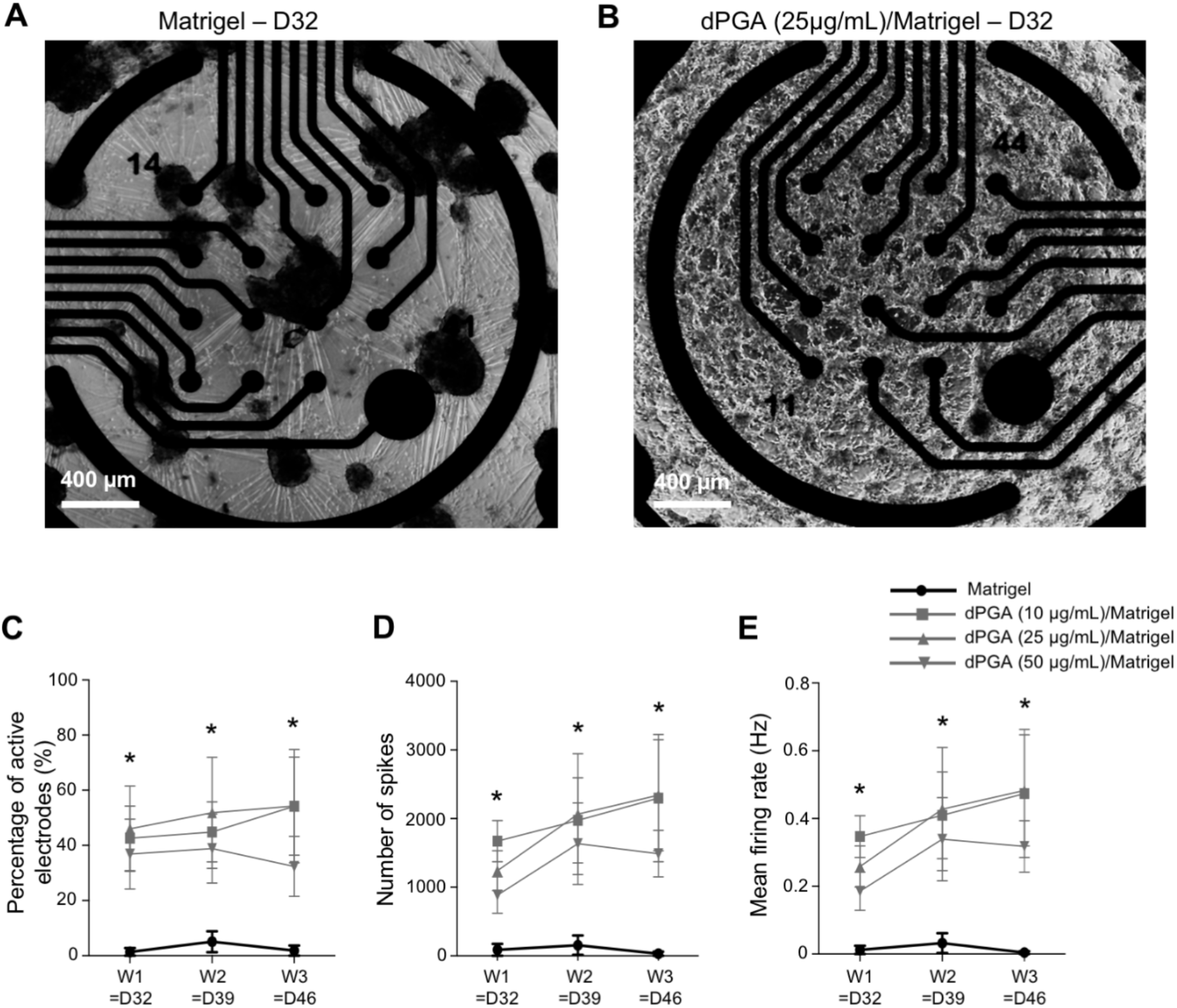
The dPGA/Matrigel substrate enables long-term multielectrode array recordings of human iPSC-derived motor neurons. **A-B)** Representative images of h-iPSC derived-motor neurons (MNs) one-week post-plating (i.e., 32 days of differentiation) on MEA wells coated with either Matrigel only **(A)** or dPGA (25 μg/mL)/Matrigel **(B)**. Scale bars, 400 μm. **C-E)** Number of active electrodes (%) **(C)**, number of spikes **(D),** and mean firing rate (Hz) **(E)** of MN cultures plated on Matrigel only vs dPGA (10, 25, or 50 μg/mL)/Matrigel at one-, two- and three-weeks post-plating (n=3 cultures/condition; 6 wells for each culture; p=0.0017 for the number of active electrodes (**C**) and p=0.0174 for the number of spikes (**D**) and mean firing rate (**E**) (Friedman test); *, p<0.05 for dPGA (25 g/mL)/Matrigel vs Matrigel only (Dunn’s multiple comparisons post-hoc test). Data are shown as mean ± SEM.

### Improved quality of samples for single-cell RNA sequencing of developmentally mature human iPSC-derived motor neurons cultured on dPGA/Matrigel

Microdroplet-based scRNAseq analysis provides a powerful tool to validate the quality and composition of h-iPSC-derived MN cultures by enabling a precise characterization of the genome-wide expression profile of individual cells (Stegle et al. 2015; Thiry et al., 2020). Performing scRNAseq analysis on induced MNs that have already begun to coalesce into large cell clusters, as they do as early as one-week post-plating on Matrigel (Fig. 2), makes the isolation of viable single cells technically challenging. To address this situation, we evaluated whether the use of dPGA/Matrigel coating, which leads to reduced MN clustering, would facilitate the application of scRNAseq to the study of h-iPSC-derived MNs. We submitted h-iPSC-derived MNs plated on either Matrigel only (“Matrigel”) or 50 μg/mL dPGA/Matrigel (“dPGA/Matrigel”) to scRNAseq at 39 days of differentiation. After transcriptomic data (FASTQ files) were acquired from ~5000 single cells for each sample, we used the Bioconductor software (Lun et al. 2016) to perform quality control (QC) steps to filter out damaged cells (Figure 4A-C). A total of 4375 cells of the “Matrigel” sample and 5857 cells of the “dPGA/Matrigel” sample passed QC. Interestingly, the median number of mapped reads per cell was lower in Matrigel as compared to dPGA/Matrigel conditions [Fig. 4A; 6369 for Matrigel (top) and 9302 for dPGA/Matrigel (bottom)]. Similarly, the median number of genes expressed per cell was lower in Matrigel as compared to dPGA/Matrigel conditions [Fig. 4B; 2685 for Matrigel (top) and 3168 for dPGA/Matrigel (bottom)]. Together, these QC metrics suggest a reduced proportion of damaged cells (i.e., the cells with lower numbers of detected genes and mapped reads) in the “dPGA/Matrigel” sample compared to “Matrigel”. In addition, the percentage of mitochondrial genes present in most cells was higher in Matrigel” compared to “dPGA/Matrigel” samples [Fig. 4C; 10.3% for Matrigel (top) and 8.7% for dPGA/Matrigel (bottom)], suggesting a high proportion of damaged cells and/or debris in both samples, but in a reduced proportion in the “dPGA/Matrigel” sample as compared to “Matrigel”.

**Figure 4:**
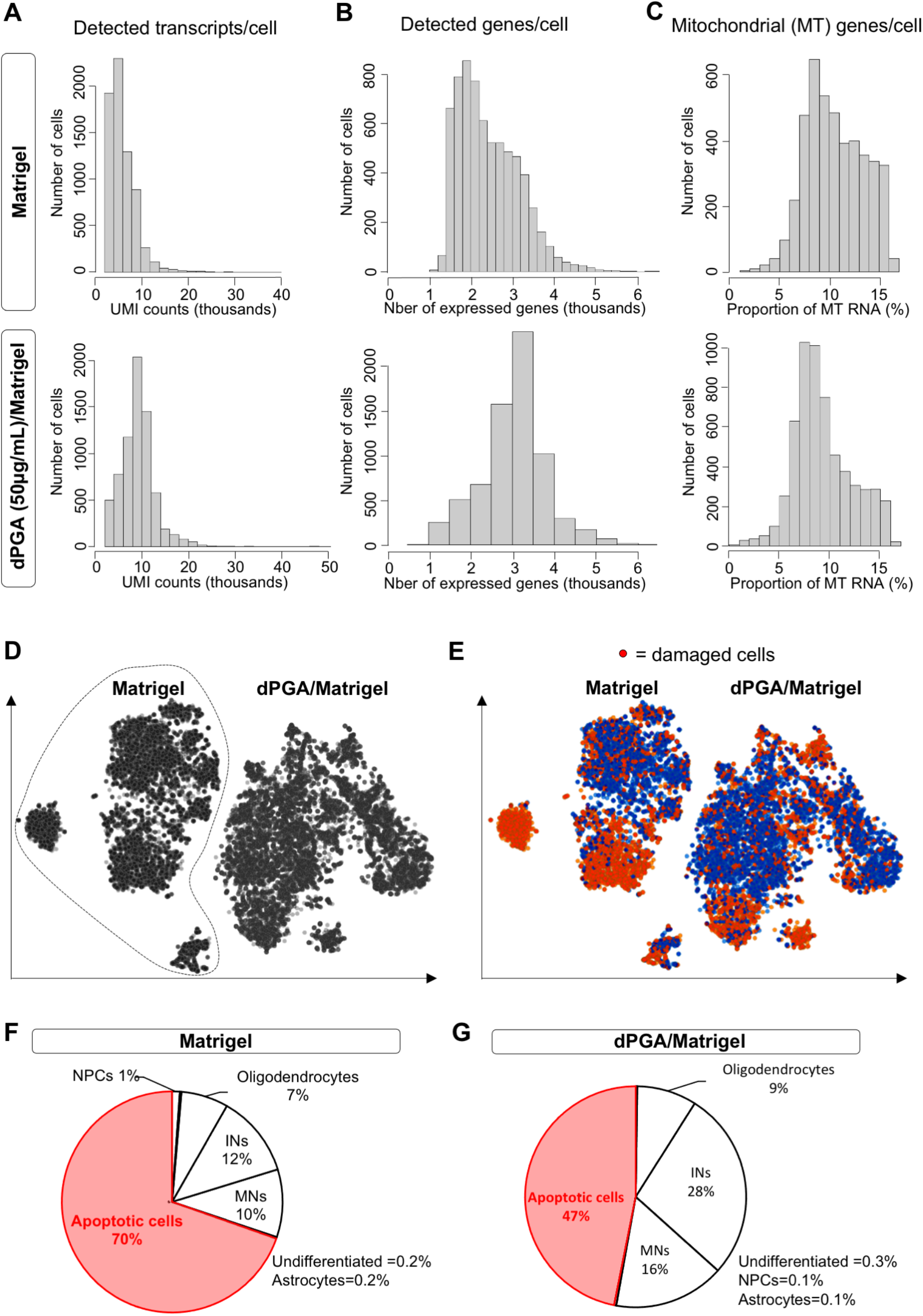
Reduced proportion of damaged cells after single cell RNA sequencing analysis of human iPSC-derived motor neuron cultures grown on dPGA/Matrigel. **A-C)** Quality control metrics for h-iPSC-derived MN cultures grown on either Matrigel only (top graphs) or dPGA (50 μg/mL)/Matrigel (bottom graphs) after 39 days of differentiation, with the number of detected transcripts per cell **(A)**, the number of detected genes per cell **(B)**, and the number of mitochondrial (MT) genes per cell **(C)** depicted. **D)** t-SNE plot of h-iPSC derived MN cultures grown on either Matrigel only (circled area) or dPGA (50 μg/mL)/Matrigel after 39 days of differentiation. Each dot corresponds to a cell. **E)** t-SNE plot of the same h-iPSC-derived MN cultures as in **D**), colored in red for cells identified as damaged (based on low QC metrics and/or expression of known apoptotic markers such as *BCL2*, *BAX*, *NGF, BOK, ATM, PSEN1, RB1, LIG4, CDK5, PARP1, BAD, MAP3K11, UBE2M)*. **F-G)** Relative proportion of undifferentiated cells, NPCs, interneurons (INs), MNs, astrocytes, oligodendrocytes, and damaged cells, identified two weeks post-plating (i.e.; 39 days of differentiation) on either Matrigel only **(F)** or dPGA (50 μg/mL)/Matrigel **(G)**, based on their level of expression of the following genes: *NANOG* and *OCT4* for undifferentiated cells; *SOX1*, *SOX2*, and *MKI67* for NPCs; *PAX2*, *PAX3*, *LBX1*, *EVX1*, *EN1*, *CHX10*, *GATA3*, *SOX14*, *SIM1*, *TLX3* for INs; *NEUROG2*, *OLIG2*, *HB9*, *ISL1*, *ISL2*, *CHAT* for MNs; *S100B and SOX9* for astrocytic glial cells; *PDGFRα* and *GALC* for oligodendrocytes; *BCL2*, *BAX*, *NGF, BOK, ATM, PSEN1, RB1, LIG4, CDK5, PARP1, BAD, MAP3K11, UBE2M* for apoptotic cells.

After QC, the remaining cells were submitted to principal-component analysis and t-distributed stochastic neighbor embedding (t-SNE) analysis (Figure 4D). In both samples, a large number of damaged/dying cells were identified, based on their low QC metrics and/or their high levels of expression of known markers of cell death such as *BCL2*, *BAX*, *NGF, BOK, ATM, PSEN1, RB1, LIG4, CDK5, PARP1, BAD, MAP3K11, UBE2M* (Wei et al. 2001, D’Orsi et al. 2015; Frade and Barde, 1999) (Figure 4E). To further characterize this situation, we next performed integrative analysis to compare the cell proportions and gene expression differences between “Matrigel” (Figure 4F) and “dPGA/Matrigel” (Figure 4G) samples. Although a large proportion of cells were identified as “damaged cells” in both samples, this fraction was markedly reduced in the “dPGA/Matrigel” sample (47%; Figure 4G) as compared to “Matrigel” (70%; Figure 4F). Consequently, a larger proportion of cells were identified either as MNs and MNPCs (“MNs”; 16% in dPGA/Matrigel vs. 10% in Matrigel) based on their expression of the specific markers NEUROGENIN2 (*NEUROG2*), *OLIG2, HB9, ISL1, ISL2* and CHOLINE ACETYL TRANSFERASE (*CHAT*), or as interneurons (“INs”; 28% in dPGA/Matrigel vs. 12% in Matrigel) based on their expression of the specific markers *PAX2*, *PAX3*, *LBX1*, *EVX1*, *EN1*, *CHX10*, *GATA3*, *SOX14*, *SIM1*, *TLX3*. Of note, other cell types were identified in relatively comparable proportions in both samples: oligodendrocytes accounted for only 7% of the cells in Matrigel and 9% in dPGA/Matrigel, while undifferentiated cells (iPSCs), NPCs and astrocytes were almost absent in both samples, as expected.

Together, these scRNAseq results show that although it is technically challenging to isolate single MNs from ‘more mature’ h-iPSC-derived cultures without causing cell damage in the process, the quality of single-cell suspensions for scRNAseq can be significantly improved by the use of dPGA/Matrigel coating during the maturation of h-iPSC-derived MN cultures.

## Discussion

Despite rapid advancements in iPSC-based technologies, current neuronal differentiation protocols tend to generate cells with properties of relatively immature neurons. This poses a problem when using iPSC-derived neurons to model age-related pathologies, including ALS (Arbab, Baars, & Geijsen, 2014; Luisier et al., 2018; Mertens et al., 2015; Miller et al., 2013; Patani et al., 2012; Ho et al. 2016). Transcriptomic comparison of h-iPSC-derived spinal MNs suggests that they are more similar to fetal than adult spinal cells (Ho et al. 2016; Luisier et al., 2018). Maturation- and age-related pathways are thought to play roles in MN disease pathology, highlighting the necessity to develop strategies to study ‘*in vitro* matured’ h-iPSC-derived MNs to provide more disease-relevant iPSC-based models of ALS. However, h-iPSC-derived MN maturation is made difficult in most current derivation protocols by the appearance of large MN aggregates, often as early as 5-7 days post-plating, which cause significant cell loss and hinder the analysis of single cells (Kuijlaars et al. 2016; Taga et al. 2019; Thiry et al. 2020). This shortcoming underscores a critical need to improve long-term culture of h-iPSC-derived MNs to increase the fractions of viable single cells. Based on these observations, the present study aimed at developing enhanced culture conditions to facilitate the study of cell viability, molecular identity, and network electrophysiological activity in long-term h-iPSC-derived MNs.

The extracellular matrix (ECM) performs a number of important functions, including mediation of cell–cell interactions, cell adhesion and migration, control of cell proliferation and differentiation, and apoptosis (Kharkar et al. 2013; Wang et al. 2015). Over the past decades, the development of artificial ECM materials, for example via the use of ECM-mimicking substrates like Matrigel, has allowed the *in vitro* expansion and differentiation of iPSCs in a controllable, robust and scalable manner (reviewed in Schmidt et al 2017). However, such protein-based substrates are susceptible to degradation by enzymes secreted by the cells, impacting on their use for long-term culture. To overcome this situation, non-peptide polymer substrates that are resistant to enzymatic degradation by the cells, like the cytocompatible but non-biodegradable dPGA (Clement et al. submitted), can be used in combination with peptide-based substrates to achieve a balance between resistance to protein degradation and lack of inherent biological activity of synthetic polymers. The present study has provided evidence suggesting that the use of this approach has the potential to improve the study of h-iPSC-derived MNs by enhancing morphological, immunocytochemical, and survival studies, extracellular electrophysiological recordings, and single-cell RNA sequencing.

Our results have shown that dPGA coating in conjunction with Matrigel does not interfere with the differentiation of h-iPSCs into MNs. More importantly, it results in a more homogeneous distribution of h-iPSC-derived MNs, with widespread arborizations of their processes after the first week post-plating. These improved conditions result in a significant increase in MN survival over time, when compared to coating without dPGA. This advancement is expected to allow improved long-term qualitative and quantitative morphological and immunocytochemical analysis, as well as survival studies, of more mature h-iPSC-derived MNs. This progress will benefit ALS research based on h-iPSC-derived MNs in a number of ways. For instance, it will facilitate the study of how different subtypes of spinal MNs, differing in gene expression, connectivity and function (William, Tanabe, & Jessell, 2003), are specifically vulnerable to degeneration in ALS (Chakkalakal, Nishimune, Ruas, Spiegelman, & Sanes, 2010). Being able to identify each subtype present in more mature iPSC-derived MN cultures, and to monitor changes in the morphology and survival rate of each MN subtype, will provide insight into the mechanisms responsible for driving differential susceptibility to degeneration. More generally, although a growing number of studies of immature MNs derived from ALS patient iPSCs have shown their potential in recapitulating early morphological and disease-relevant molecular phenotypes, including RNA foci, protein inclusions, neurofilament aggregation, nucleoplasmic transport disruption, and cell death (Sareen et al. 2013; Chen et al. 2014; Kiskinis et al. 2014; Zhang et al. 2015), it remains to be determined how these phenotypes contribute to disease progression in more mature MNs. The availability of experimentally tractable ‘*in vitro* matured’ cultures will provide better model systems to answer these questions.

Beyond morphological, immunocytochemical and survival studies, the electrophysiological characterization of h-iPSC-derived MNs is crucial to obtain accurate measures of their function. Multielectrode arrays are particularly suited for electrophysiological studies as they enable the extracellular recording of large populations of neurons, allowing for extended recordings that are important for a number of objectives, including drug testing. As shown in this study, and previously observed by others (Kuijlaars et al. 2016; Taga et al. 2019), h-iPSC-derived MNs plated on a peptide-based substrate typically form a significant number of large cell aggregates that make it difficult to discern specific synaptic connections between the cells and to record their electrophysiological activity. We have shown that the plating of h-iPSC-derived MNs on dPGA/Matrigel coated-MEA plates results in a more homogeneous distribution of the cells and a greater number of electrodes showing persistent activity overtime. Such improved experimental conditions will enable the monitoring of the electrophysiological activity of h-iPSC-derived MNs over extended periods of time, an important goal of ALS research. For instance, analysis of MNs generated from C9ORF72 patient-derived iPSCs revealed that these cells display an initial period of hyperexcitability followed by a progressive loss in action potential output and synaptic activity at later timepoints (Devlin et al. 2015). These temporal changes could explain why several studies reported divergent findings regarding the inherent excitability of ALS iPSC-derived MNs (Sareen et al. 2013; Wainger et al. 2014; Naujock et al. 2016). While electrophysiological changes are a major phenotype in iPSC-based models of ALS, whether hyper- or hypo-excitability plays a crucial role in the disease remains unclear, highlighting the need to study the electrophysiological properties of ALS iPSC-derived MNs over extended periods of time in a stable and scalable manner.

The study of h-iPSC-derived MNs using scRNAseq becomes increasingly challenging as cultures mature *in vitro*, most likely due to the harsher conditions needed to dissociate larger MN clusters. In previous studies, we showed that scRNAseq data obtained from relatively immature h-iPSC-derived MNs (28 days of differentiation) identified approximately 4% of dying cells (Thiry et al. 2020). In contrast, the present studies have shown that a significantly larger proportion of more mature cells (39 days after differentiation) were identified as damaged cells using scRNAseq. However, we have shown that the proportion of damaged cells in h-iPSC-derived MNs cultures subjected to scRNAseq at later timepoints can be significantly reduced by the use of the dPGA/Matrigel combination as a coating substrate. This finding is expected to facilitate future scRNAseq studies of more mature h-iPSC-derived MN cultures, thus enabling gene expression profiling of cells more likely to provide information on phenotypes that may be more relevant to MN disease pathophysiology.

In summary, improved experimental conditions for the long-term culture of MNs generated from h-iPSCs are expected to promote advancements in various fields of research. They will facilitate the study of more mature h-iPSC-derived MNs, by enabling a better application of cell imaging techniques, electrophysiological recordings, as well as scRNAseq approaches, all of which are heavily reliant on the quality of the long-term MN cultures. In turn, improved long-term culture of h-iPSC-derived MNs will facilitate the study of phenotypes associated with neurodegeneration, thus providing a more informative experimental model system to develop cellular assays and facilitate early-stage drug discovery efforts in ALS and other age-related MN diseases.

## Data Availability

All data underlying the results are presented here.

## Author Contributions

LT designed and performed all experiments, data analysis, figures and wrote the first draft of the manuscript. SS, LT, J-PC and TEK conceived the study plan. RH provided and validated the dPGA. SS supervised data analysis and manuscript writing.

## Acknowledgments

We thank Dr. Thomas Durcan for advice and for providing access to multielectrode array recording set-up, Dr. Ragoussis’ team at the McGill University and Genome Quebec Innovation Center for scRNA library preparation, as well as Rita Lo for invaluable assistance. We also acknowledge Cathleen Schlesener and Markus Hellmund for their help in the dPGA synthesis.

